# Feeding experiments with Senegal bichir *Polypterus senegalus* provide evidence for diet-related tooth damage and dental microwear

**DOI:** 10.64898/2026.11.01.763519

**Authors:** Tai Kubo, Johannes N. Wibisana, Ray A. Sallan, Lauren Sallan

**Affiliations:** Center for Data Science, Waseda University, Shinjyuku-ku, Tokyo, Japan; Macroevolution Unit, Okinawa Institute of Science and Technology, Onna - son, Okinawa, Japan; Science and Technology Group, Okinawa Institute of Science and Technology Graduate University, Onna-son, Okinawa, Japan

**Keywords:** feeding experiments, *Polypterus senegalus*, dental microwear, tooth damage

## Abstract

Dental microwear is a useful tool for reconstructing jaw movements and diet in toothed vertebrates, but its application to fishes (Actinopterygii and Chondrichthyes) remains limited. This is partly due to the lack of experiments directly linking diet and microwear in these groups; only one highly-derived teleost (Actinopterygii, or ray-finned fishes) species of recent origins, the three-spined stickleback Gasterosteus aculeatus, has been subject to controlled feeding experiments. As sticklebacks are suction or ram feeders that live on macroplankton and insect larvae and do not use their dentitions for prey processing, the absence of experimental data from other fish species limits our ability to interpret dental microwear and infer diet and feeding style in a wider range of fossil and living species with various ecologies. Here we present the results of a second feeding experiment involving fishes using the basally-branching, biting, often-piscivorous ray-finned fish Senegal bichir *Polypterus senegalus* to evaluate whether controlled diet differences can generate detectable tooth damage and dental microwear patterns. Individuals were fed four different food items—loaches, shrimp, pellets, and mealworms—for one, two, and three months. We then examined tooth tip damage and linear microwear (scratches). Shrimp and mealworm feeding produced short scratches, whereas pellet and loach feeding produced long scratches. Most teeth exhibited apical tip damage despite the presence of a solid enameloid acrodin cap (a characteristic of ray-finned fishes), with damage being larger in loach-fed individuals and smaller in those fed with mealworms. Our results suggest a possible relationship between diet, tooth damage, and dental microwear in non-durophagous fishes which never-the-less use teeth for prey capture and processing, offering a framework for inferring diet in extinct forms with similar ecomorphology to *Polypterus*.

## Introduction

Microscopic scars formed by tooth-tooth and tooth-food contact during oral food processing are referred to as dental microwear. The shape and pattern of microwear marks have been used as indicators of jaw movement and dietary preference in both extinct and extant organisms with uncertain feeding ecology (Schmidt and Ungar, 2023). The application of dental microwear analysis was first limited to mammals and it is still almost exclusively used in that group (Schmidt and Ungar, 2023; Ungar et al., 2008). The first application outside the lineage of mammals was on an extinct jawless vertebrates, the conodonts (Purnell, 1995). Thereafter, a small series of studies on fish dental microwear was carried out, such as on living and fossil threespine stickleback involving both feeding experiments and dietary reconstruction (Purnell et al., 2007, 2006). Other studies have used three-dimensional dental microwear texture analysis (DMTA), which utilizes surface roughness parameters calculated from the x-, y-, and z-coordinates of microscopic areas of the tooth surface, to distinguish wild populations of cichlids, seabream, and wolffish with different dietary preferences, sometimes supplemented by data from a captive population (Baines et al., 2014; Purnell et al., 2012; Purnell and Darras, 2016). Likewise, non-experimental surveys of chondrichthyans have been conducted to infer the functions of rostral teeth in extant and fossil chondrichthyans (Itano, 2019; Nevatte et al., 2017), and to test the relationship between wear patterns and diets across living shark species to infer the diet of extinct sharks (Paredes-Aliaga et al., 2024; Paredes-Aliaga and Herráiz, 2024). Further, DTMA was used to differentiate wild and captive Sand tiger shark (McLennan and Purnell, 2021) and to infer the opportunistic feeding ecology of the Devonian chondrichthyan *Ctenacanthus concinnus* (Greif et al., 2025). These studies on dental microwear of fishes generally showed that fishes fed on harder food exhibited rougher tooth surface in three-dimensional analysis and dense and short scratches in two-dimensional analysis (Purnell et al., 2006; Purnell and Darras, 2016), although Weber et al. (2021) indicated that different species of wild extant shark with different diets could not be discriminated by DMTA,

The number of dental microwear studies focusing on fish, as shown above, is much smaller than that on other vertebrates, particularly mammals (e.g., Calandra and Merceron, 2016; Desantis, 2016). In addition, only one species, the three-spined stickleback, had been subject to controlled feeding experiments that directly relate controlled intake to observed microwear, while all other observations are based on inference from general dietary ecology. This situation is surprising, especially considering that fishes account for more than half of all vertebrate species (Friedman and Sallan, 2012). The almost complete absence of direct experimental or survey data, for most species presents a number of issues for interpreting dental microwear across extinct and extant biodiversity. Because of the constraints imposed by feeding in water, most fishes primarily rely on suction or ram feeding to obtain their prey, which is different from terrestrial vertebrates (Wainwright et al., 2015). Very few species directly capture or process prey primarily through tooth contact, with notable exceptions among durophagous and other manipulative feeders (Ferry-Graham and Lauder, 2001). Because of these differing demands, tooth morphology within fishes varies significantly: from molariform teeth of durophagous taxa to conical teeth of non-durophagous taxa (e.g.; see Purnell et al., 2006; Purnell and Darras, 2016). Therefore, more investigation into the etiology of fish dental microwear is needed to understand how different feeding styles and diets interact with different tooth morphologies, and ultimately how feeding demands have affected the evolution of dental traits in fish species.

There are three main approaches to confirm the relationship between diet and dental microwear characteristics: (i) *in vitro* studies of preserved teeth, (ii) comparative studies that involve species or populations with different known dietary preferences, and (iii) *in vivo* studies (Schmidt and Ungar, 2023). To date, there have been no *in vitro* studies on fish teeth that observe dental microwear patterns formed under different loads and abrasive types using machines that simulate fish chewing. Nevertheless, an *in vitro* study on conical-shaped alligator teeth may serve as a useful reference for fish teeth with similar morphology, accounting for differences in enameloid (Usami and Kubo, 2025). There are several comparative and *in vivo* studies on fishes as noted above (McLennan and Purnell, 2021; Paredes-Aliaga and Herráiz, 2024; Purnell et al., 2012; Purnell and Darras, 2016). The only *in vivo* study on non-durophagous bony fish, Purnell et al. (2006) on sticklebacks, examined two feeding regimes (limnetic and benthic) that combined different substrates and food types. However, as noted above, other studies on fish dental microwear compared captive individuals fed controlled diets with wild populations, rather than with other captive individuals of the same species raised on different diets (McLennan and Purnell, 2021; Purnell et al., 2012; Purnell and Darras, 2016). To fill the gap in experimental studies comparing dental microwear of conspecific fish fed on different dites, we conducted a feeding experiment using Senegal bichir (*Polypterus senegalus*). *Polypterus* belong to the sister group to all other actinopterygians (ray-finned fish) (Sallan, 2014) and are often used as extant analogues of stem tetrapods (e.g., Graham et al., 2014; Standen et al., 2014). Bichirs grow to substantially larger sizes than sticklebacks (>20 cm at maturity), can feed by dental grasping, biting and other oral processing rather than suction alone, and can be piscivorous (Whitlow et al., 2022; Hickley and Bailey, 1987). These feeding and dietary traits may facilitate greater and/or different microwear damage than that observed in small, suction/ram-feeding sticklebacks fed on soft macroplankton and smaller insect larvae (Purnell et al., 2006). Therefore, feeding experiments using *Polypterus* can fill gaps in our understanding of the relationship between dental microwear features and food properties in ray-finned and other osteichthyan (bony) fishes. It may serve as a basis to understand the feeding ecology of stem-tetrapods and other early crown vertebrates with similar dental characteristics, through comparison of dental microwear.

The aim of this study is to test whether different food items induce characteristically different patterns of dental microwear in Polypterus. Also, we investigated how other factors, such as relative tooth position and body size—a factor that defines bite force (Bestwich et al., 2021)— affected dental microwear. In addition, during the course of the experiment, we found that in many teeth, part or all the solid tip enameloid (acrodin cap, an ancestral trait for ray-finned fishes) had detached antemortem (Fig. 1). We measured the size of such damage to test whether it also correlated with diet, as this may indicate susceptibility that could have influenced the evolution of this structure. For these purposes, we fed four food items to different individuals of *Polypterus senegalus* for different durations and examined how different treatments affected the lengths and densities of scratches (linear dental microwears features) on tooth surfaces.

**Figure 1.**
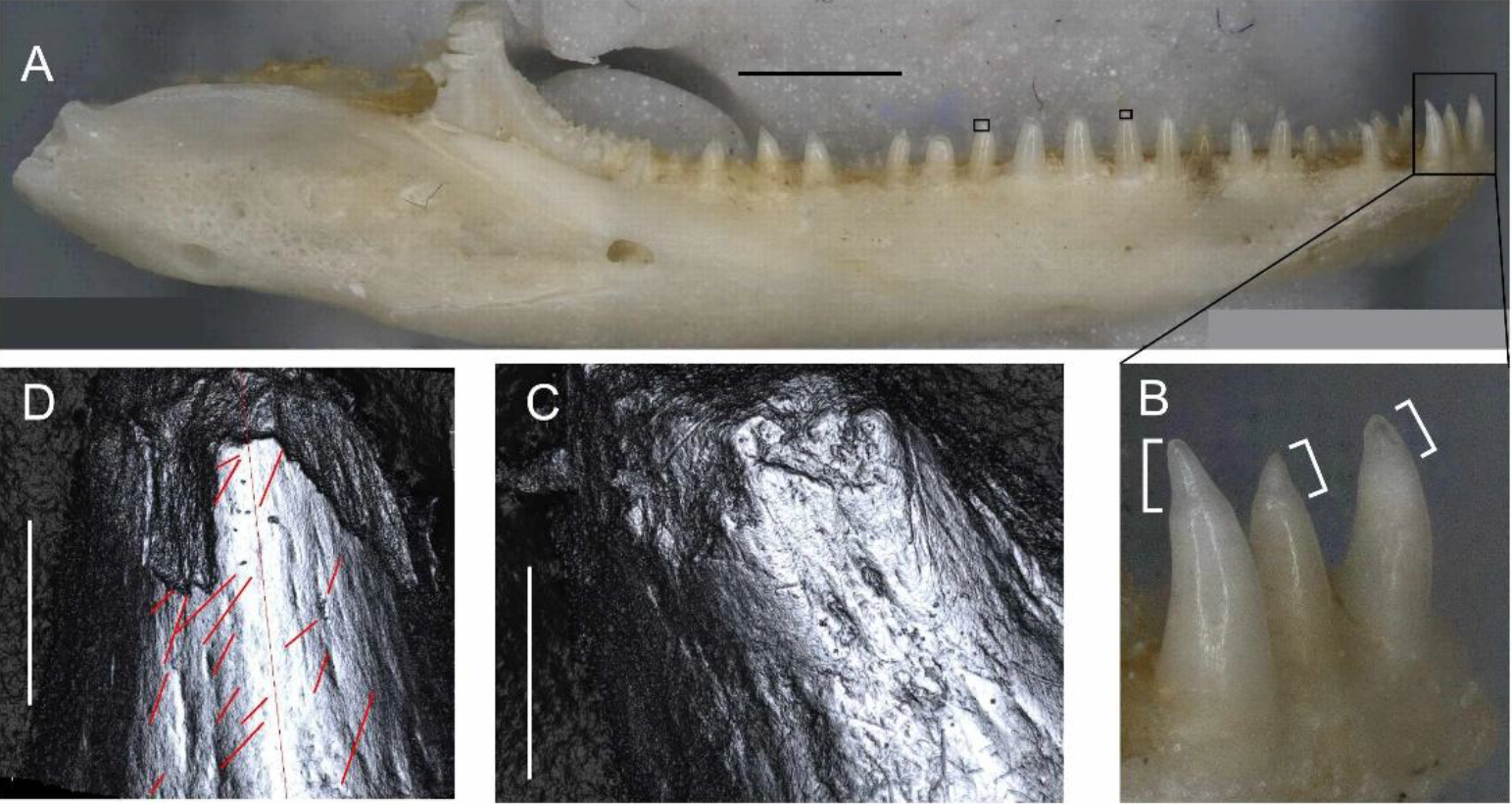
Right lower jaw and photosimulations of tooth apical tips of *Polypteru*s fed with loach for one month. (A) Buccal view of the right lower jaw. Small open black rectangles indicate the areas scanned for images C and D, from left to right. (B) Close up of mesial three teeth. Acrodin caps are highlighted with square bracket shaped symbols. (C) 10th anterior tooth, whose apical tip was broken off. Some scratches are observed on the surface that is exposed after the tip was broken off. (D) 13th tooth, showing how scratches were measured using ImageJ. Red lines were drawn where scratches were recognized. The long red line at the middle was drawn to indicate the orientation of tooth longitudinal axis. Scale bars: 2 mm in A and 50 μm in C and D.

## Material and methods

### The schedule of the experiment

We used 12 individuals of *Polypterus* with total lengths ranging from 189 mm to 238 mm. Four food items—loach, shrimp, mealworm, and pellets—were fed for durations of one, two, and three months (Fig. 2). Because there was only the room for five fish tanks with each separated into two sections containing total of 100L water, we could not conduct feeding experiments on all twelve *Polypterus* in parallel. Accordingly, we first purchased 10 individuals of *Polypterus senegalus* from an online pet shop in Japan and started feeding them pellets (Hikari Crest Freak Bottoms #4971618-274151 [Hikari, Japan]) on March 15th 2023. Feeding experiments started on November 29th 2023, after sufficient time had passed to minimize pre-existing damage. Although tooth replacement rates of ray-finned fishes have been not well studied and the exact tooth replacement rate for *Polypterus* is unknown, average tooth lifespan for Pacific lingcod is less than a month (Carr et al., 2021). So, we assume this pre-experiment feeding interval was sufficient to replace the teeth of *Polypterus*, which also exhibits regular tooth replacement and shedding (Vandenplas et al., 2014).

**Figure 2.**
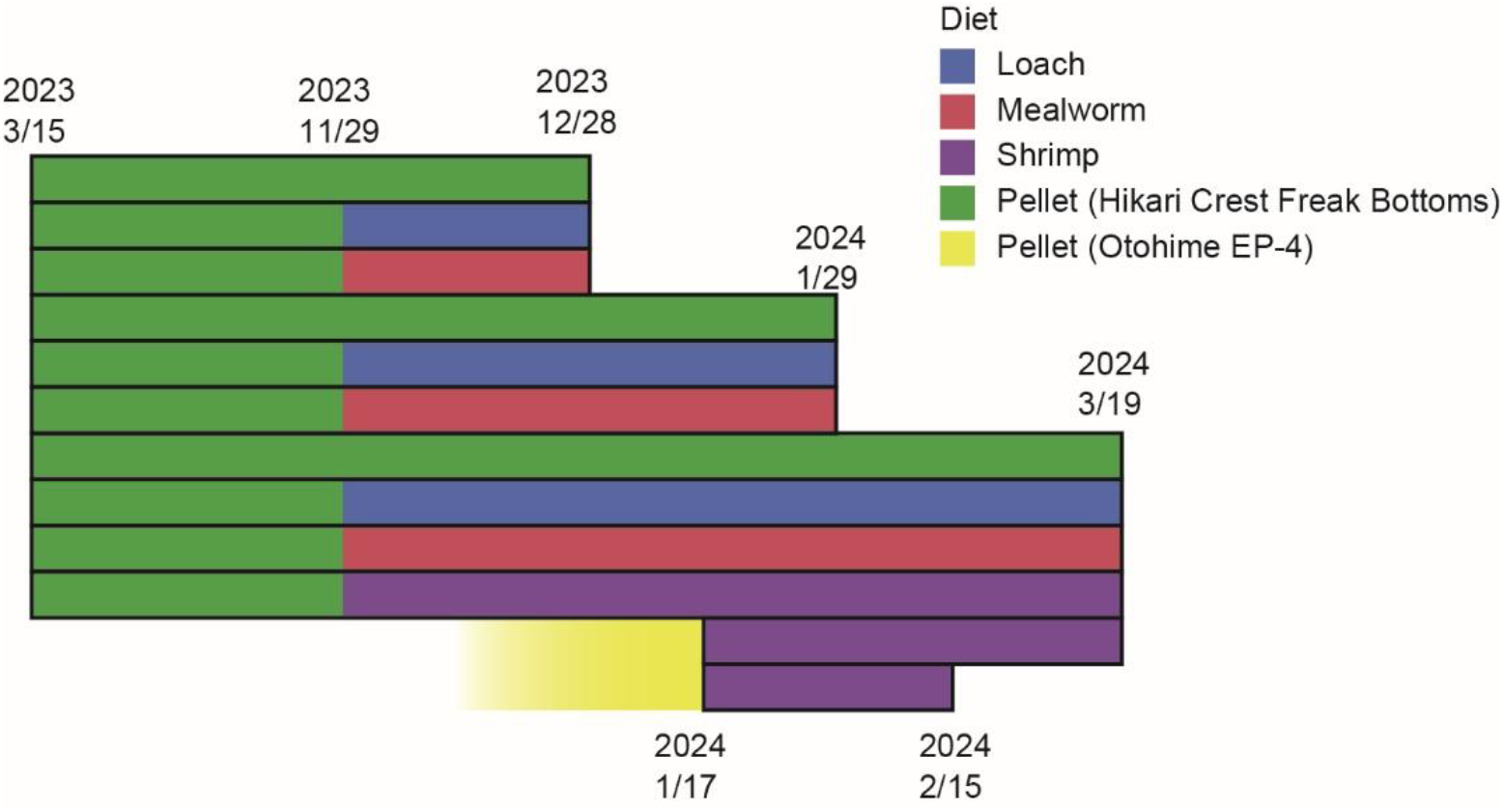
The gantt chart showing the schedule of feeding experiment for 12 individuals of Polypterus. Each box represents the feeding schedule for an individual fish.

Among these initial 10 *Polypterus*, one individual was fed exclusively shrimp for three months and others were fed exclusively pellet, loach, and mealworm for one, two, and three months. Once three individuals were euthanized from the pellet, loach, and mealworm groups for one-month sampling on 28th December 2023, an additional two *Polypterus senegalus* were puchased from an online pet shop in Japan. These two were designated to be fed exclusively shrimp for one and two months, completing the shrimp experimental set, and feeding started on 17th January 2024. These two additional *Polypterus* were reared before purchase by the pet supplier and fed exclusively pellets (Otohime EP-4 [Nisshin Marubeni, Japan]), but these were smaller and may have slightly different material properties that what we used for our initial 10 *Polypterus* specimens during the first phase of the experiment. Three individuals fed with pellet, mealworm, and loach diets for two months were euthanized on 29 January 2024. The individual fed shrimp for one month was euthanized on 15 February 2024, and all remaining five individuals were euthanized on 19 March 2024.

For euthanasia, anesthetic overdose (a 1:2 ratio of mass of tricaine methanesulfonate [MS-222 : Sigma-Aldrich, USA] and NaHCO3 [Nacalai Tesque, Japan] as a buffer) was used under the supervision of a technician from the Animal Care and Use Committee of Okinawa Institute of Science and Technology. Each fish was transferred to a smaller tank of 2 L of house water with 3-6 % dissolved buffered MS-222. Death was confirmed by the complete cessation of opercular movement for several minutes. This method is widely used for fish euthanasia and has also been applied to *Polypterus* (Lutek and Standen, 2021). In Japan, only experiments using tetrapods require formal governmental approval, while experiments on fish are approved at the institutional level. Therefore, all procedures were approved prior to the start of the experiment by the ethics committee of the Okinawa Institute of Science and Technology under protocol number ACUP-2023-006-2.

### Conditions of feeding experiment

We followed the protocol of Wibisana et al. (2025) for the husbandry of *Polypterus senegalus*. Each *Polypterus* was kept in a separate section and fed a single food item three times per week: two pellets (Hikari Crest Freak Bottoms #4971618-274151 [Hikari, Japan]), two live mealworms (*Tenebrio molitor*, about 3 cm in length), two small live loaches (*Misgurnus anguillicaudatus*, about 7–10 cm in length), or two frozen shrimps (*Caridina multidentata* and *Neocaridina denticulata*, about 2–4 cm). Shrimps were not defrosted before the feeding. When possible, videos of *Polypterus* biting food items were recorded using the feeder’s smartphone for bite counting.

Water temperature and pH were recorded at each feeding. The water temperature was kept to be 25 degrees Cersius. For water conditioning, Eheim Classic 2215 external filters (Eheim,Germany) were used and 10-30 % of the water was replaced every week. Any leftover food was removed, except for loaches. When loaches remained, we adjusted the number of loaches at the next feeding schedule so that the total number was two.

### Collecting data of dental microwear

After euthanasia, all specimens were preserved as wet specimens using 10% formalin and 70% ethanol following the steps 12–14 of the protocol established by Motomura and Ishikawa (2012). For dental microwear analysis, the section of left lower jaw (dentary) bearing marginal teeth was removed from each specimen. When possible, soft tissues were physically removed, and the tooth surfaces were cleaned with acetone–soaked cotton swabs (Fig. 2). If the tooth surfaces were still covered by organic material afterward, they were further cleaned in an ultrasound bath. Then, jaws were individually placed in a 2% NaClO solution (sodium hypochlorite bleach) for 1 minute, transferred to a water bath for 1 minute, and cleaned again with acetone-soaked cotton swabs. The weight of each wet specimen (drained in ethanol) was measured together with the corresponding left lower jaw specimen to be used as a body size index.

The buccal side of teeth in the largest and most buccal marginal row was scanned using a confocal laser microscope (VK-X3000, Keyence, Osaka, Japan) with a violet laser (404 nm), equipped with a long working distance 100x lens (N.A. = 0.73, resolution in x, y = 0.137 μm, step size in z = 0.1 μm). Scans were conducted for all teeth preserved in left lower jaws and were assigned serial numbers (Tooth-1, Tooth-2, Tooth-3, …) from mesial to distal. For observed 12 specimens, the total number of teeth ranged from 19 to 29. For each tooth, scanning areas were designated sequentially (1, 2, 3) from the apical toward the basal direction, regardless of whether the apical portion was missing due to antemortem damage. All scans were taken so that there was no overlap.

Because the teeth were small and their buccal surfaces strongly curved, three-dimensional surfaces obtained by scans contained too much noise and could not be used for analysis. Therefore, we analyzed two-dimensional images obtained by photosimulation of the three-dimensional surface models. Although some areas showed enameloid flaking, no pit-like dental microwear was observed. Accordingly, we measured only scratches (linear dental microwear). Only scratches on the enameloid of the acrodin cap were measured. From each image, the lengths of scratches were measured using the software Image J (version 1.53e, National Institutes of Health, USA). When an image was too blurred (i.e. too much noise in the original 3D scan) or remained covered by organic material after the above treatment, no measurements were taken from that image. When scratches extended beyond the image or beyond the acrodin cap, only the length within the image or acrodin cap was measured. For curved scratches, the length was measured by connecting the two end points. Images were usually not filled with tooth surface and areas close to the edge of tooth were too steep to recognize scratches, so to standardize scratch counts, the area in which scratches could be recognized were measured for each image. These measurements were conducted solely by Tai Kubo to avoid inter-observer error from 27 June to 12 September 2025. No formal repeat analyses were performed to assess potential observer drift, due to the difficulty of training.

In addition, we found that the apical tip of many teeth had dislodged during life. This damage was judged as antemortem because scratches were sometimes present on the surface exposed after tooth breakage and loss of solid acrodin (Fig. 1), and were also potentially produced during food processing after the loss of the cap and collar enameloid on areas adjacent to the acrodin cap. We quantified the size of this damage by measuring the width of the base of the missing apical tip, defined as the length perpendicular to the longitudinal axis of the tooth. As the image width of the photosimulation is 140 μm, when this width exceeded 140 μm, it was recorded as >140 μm. For the non-parametric analysis, the value of >140 μm was treated as 140 μm, indicating that they were larger than all other values, while no order was assumed among values above 140 μm.

### Statistical analyses of dental microwear and tooth tip damage

To identify factors influencing the characteristics of dental microwear, analyses were conducted on metrics representing scratch length and scratch count per tooth. These two parameters were treated as response variables. Scratch count per tooth was analyzed using the generalized linear model as its distribution is non-normal. The total surface area per tooth (μm^2^) was incorporated as offset in this generalized linear model, which makes the analysis identical to analyzing scratch density (scratch count /μm^2^). As scratch length exhibited a strongly right-skewed distribution, it was log-transformed to approximate normality to be analyzed using the linear model.

The candidates of explanatory variables were different for scratch length and scratch count, because scratch count was collected for each tooth, whereas scratch length can be measured for each scratch. For the analyses of scratch length, candidates of explanatory variables are: diet (pellet, shrimp, mealworm, loach), reared month (1, 2, 3), apico-basal position within the tooth, relative tooth position (serial tooth number divided by entire tooth number of each individual), body weight (g), and the interaction between diet and reared month. For the analysis of scratch count per tooth, candidates for the explanatory variables were: diet, month, relative tooth position, body weight, and the interaction between diet and month. Diet was treated as a categorical variable, and all other variables were treated as continuous. When no scratches were counted on a tooth, two different treatments were applied depending on the condition of the images. If the images were clear but no scratches were observed, the tooth was recorded with a scratch count of zero. However, if the scratch count was zero because all images of that tooth were blurred, the data for that tooth were not included and instead were treated as missing values.

To identify the most relevant explanatory variables, we applied stepwise model selection based on the Bayesian Information Criterion (BIC). BIC penalizes model complexity, more specifically number of explanatory variables and sample size, and thus favors parsimonious models, reducing the risk of overfitting when multiple candidate predictors are considered. After selecting the optimal model, standard multiple regression was performed using the retained variables.

The model for scratch length was a multiple linear regression, as log-transformed scratch length could be assumed to follow a normal distribution. In contrast, count data are known to follow non-normal distributions, particularly when counts are small, as in the present study. Therefore, a generalized linear model with a Poisson error distribution was applied to analyze scratch counts, because the Poisson distribution is widely accepted for modeling count data. Because significant overdispersion was detected in the residuals of this analysis that means assumption of Poisson error distribution for scratch count is violated, we further analyzed by assuming more conservative negative binomial distribution for scratch count using the same explanatory variables and BIC-based stepwise model selection procedure. Explanatory variables included in the best-fitting models were interpreted as those significantly affecting the dental microwear features of *Polypterus*.

In addition, the relationship between mean scratch length per tooth and scratch density was examined using reduced major axis (RMA) regression. Teeth without scratches were excluded from the analysis. RMA was chosen as there is no assumed causal relationship between these two variables. Further, to examine whether scratch density influences the variance of scratch length, we fitted a loglinear variance model, with scratch count/area as a predictor of both the mean and variance of average scratch length per tooth. The significance of the variance component was assessed using a likelihood ratio test.

We also tested whether the size of the damage on apical tip of tooth was correlated with diet. Because there are upper (140μm) and lower (0μm) limits for the measured damage size, a non-parametric Steel-Dwass test was conducted to determine if there were any differences in the size of the lost tooth tip among Polypterus fed different diets.

For analyses on scratch length and damage sizes on tooth tip, we conducted additional analysis without data from *Polypterus* that was fed shrimp for one and two months to account for different pre-experiment condition.

All variable values are provided in Supplementary File 1. Statistical analyses for scratch lengths and the damages on apical tip were performed using JMP Student Edition version 19.0.1 (SAS Institute Inc., Cary, NC, USA). The analysis for scratch count and the reduced major axis regression between scratch density and mean scratch length per tooth were performed in the software R (version 4.5.2) with the code written in the Supplementary file 2 using data in the Scratch count tab in the Supplementary File 1.

### The analysis of bite count

Videos of feeding behavior were analyzed to count the number of bites for each food item. Recording typically began when *Polypterus* grasped a food item with its mouth and, in some cases, may have ended before the completion of swallowing, because the mouth movements for suctioning can continue for a while after the food item is completely inside the mouth and dental margins. Therefore, the recorded bite counts represent minimum estimates. A bite was defined as any nodding motion, lateral head movement, or observable movement of the food item caused by jaw action. The total number of bites was counted for each food item (Supplementary File 1). The number of videos differed among food items because behavioral responses and the time from placement to feeding varied. Feeding on loaches was rarely recorded because they were provided as free-swimming prey. Pellets were recorded less frequently because they were treated as a control food at the time of feeding. This imbalance in sample size likely reduced the statistical power to detect differences in bite counts. A non-parametric Steel-Dwass test was conducted to determine if there were any differences in the number of bites depending on food items using JMP Student Edition version 19.0.1.

## Results

For log-transformed scratch length, diet, rearing month (1, 2, 3), and apico-basal position within the tooth were selected as explanatory variables in the model with the lowest BIC (5691.8). The P-values for all these variables were less than 0.0001. The estimated coefficients were 0.227 for apico-basal position (indicating larger values toward the basal part of the tooth), 0.093 for month, and for food items −0.060 for shrimp and mealworm, and +0.060 for pellet and loach. These coefficients indicate that scratches are longer in the basal region of the tooth and, unexpectedly, increase with longer rearing periods. Scratches are also longer in individuals fed loach and shrimp than in those fed shrimp and mealworms.

When data from individuals added later that had been reared on shrimp for one and two months were excluded, owing to their different pre-experimental diets, rearing month and apico-basal position within the tooth were selected as explanatory variables with the BIC value of 4408, which showed that the effect of diet was not detected when these individuals were excluded.

For scratch count per tooth, when Poisson distribution is assumed for scratch count with area as an offset, diet, rearing month, relative tooth position, the interaction between diet and month were selected as explanatory variables in the model with the lowest BIC (2355.4). However, the dispersion parameter is 7.22, which indicates the assumption of the Poisson distribution of scratch count is violated. Therefore, we conducted the model selection procedure with the same set of variables and offset but with the assumption of negative binomial distribution for scratch count. The best model included only relative tooth position and rearing month as the explanatory variable with BIC of 1487.8. The estimated coefficients were -0.89032 for relative tooth position and - 0.32746 for rearing months, indicating scratch density decreases in posterior teeth and also with longer rearing periods.

No significant relationship was detected between mean scratch length per tooth and scratch density using RMA (P-value = 0.33). The loglinear variance model also showed that scratch density did not significantly predict mean scratch length (β = −254.49, F_1_,_5 8_._8_ = 0.25, p = 0.62), however, it revealed that scratch density was a highly significant predictor of the variance in scratch length (χ^2^ = 18.26, df = 1, p < 0.0001). The variance parameter estimate was negative (β = −529.82, 95% CI: −730.14 to −305.08), indicating that samples with higher scratch density exhibited substantially less variation in scratch length than samples with lower scratch density.

Damage at the apical tip of the tooth was commonly observed. Among the 268 teeth examined, 162 showed some form of damage, ranging from flaking of the enameloid tip to complete loss of the acrodin cap. Among six pairs of *Polypterus* diet groups, the size of this damage differed significantly between the group fed loaches and each of the other three groups (p < 0.001). It also differed significantly between the shrimp-fed and mealworm-fed groups (p = 0.0355). For other two pairs, between shrimp-fed and pellet-fed groups and between mealworm-fed and pellet-fed groups, differences were not significant (p > 0.05). The size of damage on the apical tip of tooth was large in the loach-fed group and small in the mealworm-fed group (Fig. 3). When the data taken from the late-added individuals that were fed shrimp for one and two months were excluded given the difference in diet prior to the experimental period (see above), loach feds were significantly different from the all-other diet groups (p < 0.001), but there was no difference between other diet groups.

**Figure 3.**
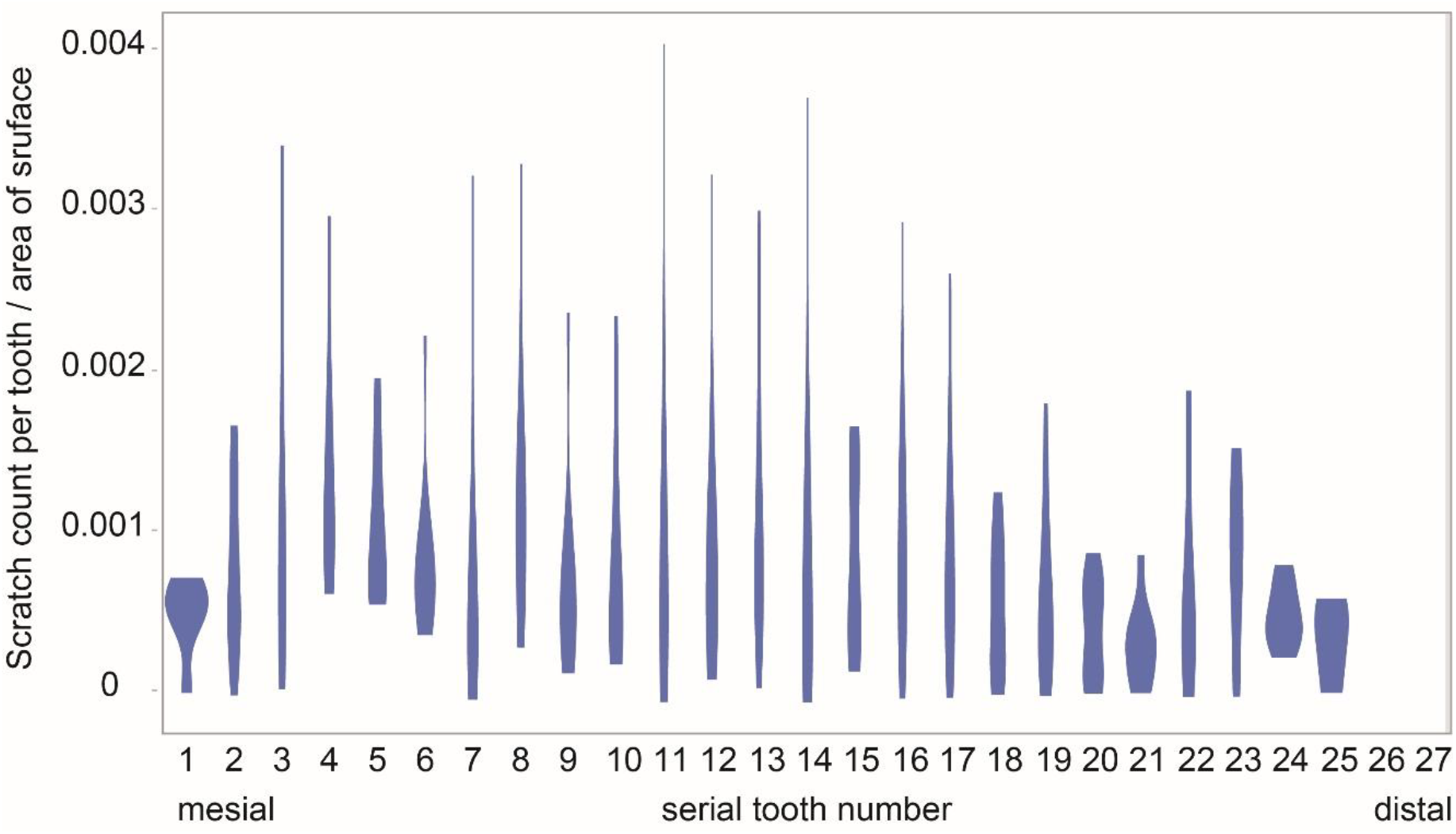
Box plot showing the size of apical tip damage on teeth. Each box represents a different *Polypterus* individual. The size of damage was defined as the width of the base of the missing apical tip. It was larger in the loach-fed group and smaller in the mealworm-fed group.

Bite counts were obtained from 23 videos recording feeding behavior involving 25 total food items: two loaches, six mealworms, one pellet, and 16 shrimps (Supplementary File 1). The status of loach as free swimming prey may have influenced biting behavior - in one video of loach feeding, no biting was observed because *Polypterus* simply held the loach for 21 seconds without biting motions. The Steel–Dwass test detected no significant differences in bite counts among the different food items (p >0.05).

## Discussion

The size of damage on the tooth tip of *Polypterus* reflected diet. We found that damage at the apical tip was severe on loach-fed individuals and less so in mealworm-fed individuals (Fig. 3). Biting counts observations did not show significant difference between different food items. However, there were only two videos for loach feeding. One video showed only four bites after capture of the loach and before swallowing that is less than any other feeding video. Another video started when Polypterus was already holding the loach and showed no additional bites; during this time, *Polypterus* held the loach for 21 seconds, which was longer than the processing time of any other food item observed. As loaches are much larger than other food items and firm, continuous contact between the teeth and body were required in at least some cases to stop movement and prevent escape, it is not surprising that tooth tips were damaged during this process. For the other food items, manipulation and capture were not as difficult: dead shrimp were fed after being frozen, and live mealworms are almost immobile in the water, so most damage was likely due to brief food-tooth contacts during biting and suction.

Diet affected the length of scratches but not the density of scratches, whereas the feeding period (months) affected both length and density of scratches. In addition, the scratch length was affected by position along the tooth surface, with scratches being longer in more basal positions. This is not surprising given the conical shape of the tooth and the fact that tooth surface becomes flatter basally. The relative position of the tooth within the row did not affect the scratch length but it affected scratch density. However, this relationship is not linear, scratch counts were low in both, the most mesial area and the most distal area (Fig. 4). This pattern may reflect that the anteroposteriorly middle teeth contact food more often during biting, as we observed that *Polypterus* manipulates prey using these teeth. Except for pellets, all food items used in this study—loaches, shrimps, and mealworms—are elongated in shape. *Polypterus* initially grasps the food by biting the anteroposteriorly middle portion of the food using its middle teeth and continues biting with these teeth along with suctioning until the food becomes aligned with the longitudinal axis of the body, allowing it to be swallowed (https://doi.org/10.5281/zenodo.20300816). Lauder (1980), reported the same feeding process in *Polypterus*: an initial strike followed by mastication involving repeated cycles of ejection and crushing, and finally rapid suction when the prey reaches the front of the jaws. Therefore, it is reasonable that middle teeth exhibit denser scratches as they were used in biting more often. Among the variables included in the model, body size and the interaction between diet and feeding period did not affect scratch length nor scratch density.

**Figure 4.**
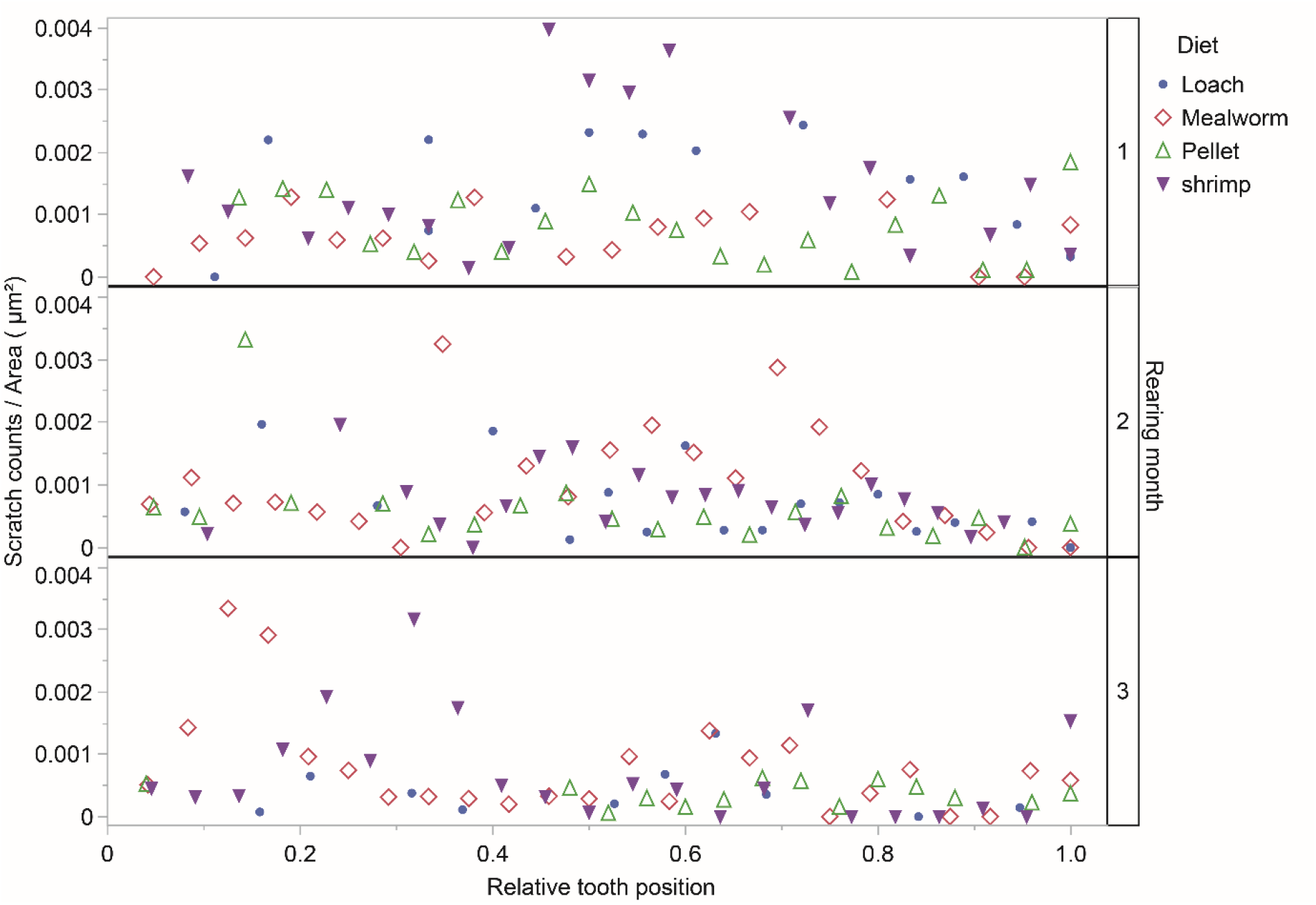
Scatter plots showing the scratch density (scratch count divided by tooth surface area) against relative tooth position (serial tooth number divided by total tooth number). Serial tooth numbers was assigned from mesial to distal, starting at 1. The number of teeth varied among individuals from 19 to 29. Scatter plots exhibiting values for *Polypterus* with feeding duration of one, two and three months from top to bottom. Scratch counts were low in the most mesial and the most distal areas but high in the middle.

Loach and pellet feeding produced longer scratches, and shrimp and mealworm feeding left shorter ones (Fig. 5), while we could not detect effect of diet on scratch density. When the relationship between mean scratch length and scratch density was examined per tooth regardless of diet in our sample, scratch density did not affect mean scratch length (p=0.33 and 0.62 for RMA and loglinear variance model respectively), but teeth with low scratch density are highly variable in their mean scratch length, whereas teeth with high scratch density are less variable (loglinear variance model: p < 0.0001). In sticklebacks, benthic populations reared in coarser substrate sediments showed shorter and denser scratches in line with their exposure to hard particles, whereas limnetic population without exposure to grit showed longer scratches with lower density (Purnell et al., 2006). Among extant elasmobranchs, durophagous taxa show a greater number of shorter scratches compared to generalist and piscivore taxa (Paredes-Aliaga et al., 2024). These two-dimensional fish dental microwear analyses reported previously may be consistent with our results in that harder food, in our case shrimp and mealworm with chitin exoskeleton left longer scratches. The effect of freezing on shrimp was not clear; however, the exoskeleton may have been thawed before being bitten by *Polypterus*, considering the small size of the shrimp and the fact that the shrimp were not immediately bitten after being introduced into the water. Nevertheless, more samples and correlation with material properties of food items are required to confirm this conclusion.

**Figure 5.**
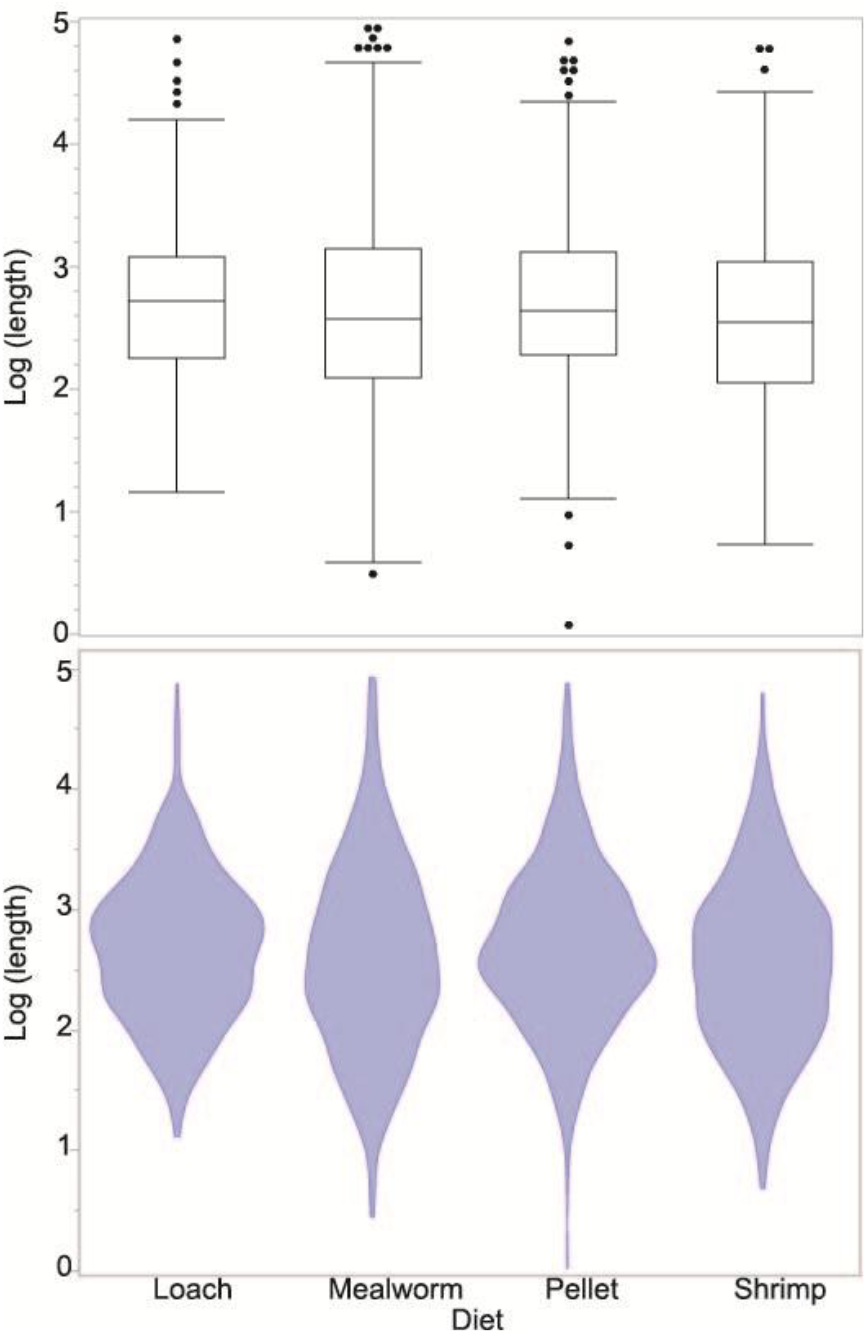
Box plot (above) and violine plot (below) showing Log length of scratches against diet. Scratches are shorter for shrimp-fed and mealworm-fed individuals, and longer for pellet-fed and loach-fed individuals.

The effect of longer feeding period (months) observed in our study is not easy to interpret. There was an effect of exposure time independent of food on microwear parameters: increasing scratch length and decreasing scratch density. Dental microwear in sticklebacks has been shown to change within four days after a diet and environmental shift (Baines et al., 2014). As we observed teeth that were one, two and three months after the dietary shift, the observed temporal change is too slow to be attributed to the effects of this shift. These temporal changes may be reflecting adaptation in feeding mode to each food item in animals exclusively fed such diets, however, there seems no clear change in biting counts over time (see Bite counts tab in the Supplementary File 1). Our sample size may be too small to reach reliable conclusions because each combination of food and duration is represented solely by single individual due to both the available room for rearing and the constraints on animal use imposed by our Institutional Animal Care and Use Committee that made it difficult to exclude the influence of individual variation.

One unexpected result was that the pellet we used left dental microwear comparable to that from other food items. We had planned the pellet as a control food causing minimal microwear, as in a previous feeding experiment on alligators (Winkler et al., 2022). This situation complicated our results and made them hard to interpret. Also, it raises the possibility that two types of pellets used for pre-experiment foods in the initial and later sets of experimental animals may have left different dental microwear. When the data from two shrimp-fed individuals that were fed different sized pre-experimental pellets were excluded, diet still affected the size of damage on the tooth tip but longer affected scratch length significantly. We note here that, for future feeding experiments; control food should be selected carefully to minimize dental microwear and pre-experimental condition should be strictly the same for all individuals.

We conducted a feeding experiment using *Polypterus senegalus* to determine the influence of diet on dental microwear and other tooth damage when controlling for other parameters. Twelve individuals were divided into four groups which were exclusively fed one of four food items: pellet, loach, shrimp, and mealworm. They were euthanized after feeding for one, two, and three months, and their dental microwear was analyzed. Unexpectedly, many apical tooth tips showed damaged (162 out of 268), all involving the solid enameloid acrodin cap which is a diagnostic feature of ray-finned fishes and represents the hardest tissue in these animals. Although this kind of damage has not been reported previously in extant fish, similar damage was reported in conodonts (Purnell and Jones, 2012), sea lion (Labrada-Martagón et al., 2007), beluga (Purnell et al., 2017), and killer whale (Foote et al., 2009), and surveys of the literature show that prior morphological studies of preserved *Polypterus* teeth likewise show this kind of wear even though it was not reported as such (Germain and Meunier, 2019: Fig. 3d). The size of the damaged or lost tooth tip reflected differences in diet. Loach-fed individuals showed the most severe damage, while mealworm fed individuals showed the least on average. Although it may be difficult to determine whether similar damage in fossil specimens occurred ante- or postmortem, the presence of scratches on surface exposed by damage can be considered as evidence of antemortem damage. It has been reported that similar tooth damage (tooth chipping) can be used as an indicator of hard-object feeding in anthropology (Towle and Loch, 2021) and as an indicator of bone consumption among large carnivorans (Van Valkenburgh, 2009; Van Valkenburgh et al., 2019). Differences in *Polypterus* feeding behavior on live versus immobile prey may be reflected in the amount and type of damage. Therefore, fish tooth-tip damage may provide a new method to infer the diet of extinct fish taxa. In addition, as *Polypterus senegalus* belongs to the sister group to all other ray-finned fishes and retains ancestral traits in tooth form, the extent of apical damage may provide some clues to as to the evolution of the acrodin cap itself.

Scratch length on the tooth surface also reflected diet, whereas scratch density did not. Among the four food items, shrimp and mealworm left short scratches. Further experimental study may be able to confirm that short scratches are characteristic for hard food, such as animals with exoskeleton. We also found that scratch density is higher in teeth in the middle of the row, that may reflect the observed feeding styles per video evidence from our experiments. Studies that link dental microwear to feeding kinematics would provide new methods to reconstruct behavior of extinct organisms.

Our experiment also provided several useful insights for future experimental design involving other fishes. A control food to establish individuals used as a baseline comparator should be selected carefully, so that it forms much less dental wear compared to other food items used in the experiment, which effect is investigated. In addition, because sample size affects statistical power, rearing a sufficient number of individuals for each food type would improve statistical robustness, although this must be balanced against the need to minimize the number of animals euthanized. When multiple individuals are kept together, it is difficult to control the amount of food consumed by each individual, however obtaining larger sample size is more important than measuring the amount of food intake for each individual.

## Acknowledgements

We thank Chen Chi Wang, Shiang Jyi Hwang and the other members of the OIST Animal Resource Section, for their assistance in fish husbandry and euthanasia. We are also grateful to all members of the Macroevolution Unit at OIST for their help with *Polypterus* husbandry and specimen preparation. We thank Mugino O. Kubo (The University of Tokyo) for allowing us to use the laser confocal microscope under her supervision, for maintaining *Polypterus* specimens, and for valuable discussions. We also thank Keita Koeda (The University of the Ryukyus) and Kotaro Shirai (The University of Tokyo) for their advice on husbandry and water sampling.

## Author contributions

Tai Kubo: Conceptualization, Methodology, Formal analysis, Investigation, Writing - Original Draft. Johannes N. Wibisana: Methodology, Investigation, Writing-Reviewing and Editing. Ray A. Sallan: Methodology, Writing-Reviewing and Editing. Lauren Sallan: Conceptualization, Resources, Supervision, Writing-Reviewing and Editing.

## Data, scripts, code, and supplementary information availability

All supplementary files, the videos used for bite counts, and the images used for the dental wear analysis are available on Zenodo (https://doi.org/10.5281/zenodo.23788078). The supplementary files include Supplementary File 1, values of all variables used in the statistical analyses and the schedule of the feeding experiments; Supplementary File 2, R code for the scratch density analysis; and Supplementary File 3, results of the statistical analyses.

## References

Baines, D. C., Purnell, M. A. and Hart, P. J. B. (2014). Tooth microwear formation rate in Gasterosteus aculeatus. J. Fish Biol. 84, 1582–1589. 10.1111/jfb.12358

Bestwick, J., Unwin, D. M., Henderson, D. M. and Purnell, M. A. (2021). Dental microwear texture analysis along reptile tooth rows: complex variation with non-dietary variables. R. Soc. Open Sci. 8(2). 10.1098/rsos.201754

Calandra, I. and Merceron, G. (2016). Dental microwear texture analysis in mammalian ecology. Mamm. Rev. 46, 215–228. 10.1111/MAM.12063/SUPPINFO

Carr, E. M., Summers, A. P. and Cohen, K. E. (2021). The moment of tooth: Rate, fate and pattern of Pacific lingcod dentition revealed by pulse-chase. Proc. R. Soc. B 288. 10.1098/RSPB.2021.1436/86400

Desantis, L. R. G. (2016). Dental microwear textures: reconstructing diets of fossil mammals. Surf. Topogr. 4, 023002. 10.1088/2051-672X/4/2/023002

Ferry-Graham, L. A. and Lauder, G. V. (2001). Aquatic prey capture in ray-finned fishes: A century of progress and new directions. J. Morphol. 248, 99–119. 10.1002/jmor.1023

Foote, A. D., Newton, J., Piertney, S. B., Willerslev, E. and Gilbert, M. T. P. (2009). Ecological, morphological and genetic divergence of sympatric North Atlantic killer whale populations. Mol. Ecol. 18, 5207–5217. 10.1111/J.1365-294X.2009.04407.X

Friedman, M. and Sallan, L. C. (2012). Five hundred million years of extinction and recovery: A phanerozoic survey of large-scale diversity patterns in fishes. Palaeontology 55, 707–742. 10.1111/j.1475-4983.2012.01165.x

Germain, D. and Meunier, F. J. (2019). Teeth of extant Polypteridae and Amiidae have plicidentine organization. Acta Zoologica 100, 119–125. 10.1111/AZO.12237

Graham, J. B., Wegner, N. C., Miller, L. A., Jew, C. J., Lai, N. C., Berquist, R. M., Frank, L. R. and Long, J. A. (2014). Spiracular air breathing in polypterid fishes and its implications for aerial respiration in stem tetrapods. Nat. Commun. 5, 1–6. 10.1038/ncomms4022

Greif, M., Calandra, I., Lautenschlager, S., Kaiser, T. M., Mezane, M. and Klug, C. (2025). Reconstruction of feeding behaviour and diet in Devonian ctenacanth chondrichthyans using dental microwear texture and finite element analyses. R. Soc. Open Sci. 12, 240936. 10.1098/rsos.240936

Hickley, P. and R. G. Bailey. (1987) Food and feeding relationships of fish in the Sudd swamps (River Nile, southern Sudan). Journal of Fish Biology 30, 147–159.

Itano, W. M. (2019). Oriented microwear on a tooth of Edestus minor (chondrichthyes, eugeneodontiformes): implications for dental function. Palaeontologia Electronica 22, 1–16. 10.26879/831

Labrada-Martagón, V., Aurioles-Gamboa, D. and Castro-González, M. I. (2007). Relation of dental wear to the concentrations of essential minerals in teeth of the California sea lion Zalophus californianus californianus. Biol. Trace Elem. Res. 115, 107–126. 10.1007/BF02686023/METRICS

Lauder, G. V. (1980). Evolution of the feeding mechanism in primitive actinopterygian fishes: a functional anatomical analysis of Polypterus, Lepisosteus, and Amia. J. Morphol. 163, 283–317. 10.1002/jmor.1051630305

Lutek, K. and Standen, E. M. (2021). Increasing viscosity helps explain locomotor control in swimming Polypterus senegalus. Integr. Org. Biol. 3, obab024. 10.1093/iob/obab024

McLennan, L. J. and Purnell, M. A. (2021). Dental microwear texture analysis as a tool for dietary discrimination in elasmobranchs. Sci. Rep. 11, 2444. 10.1038/s41598-021-81258-9

Motomura, H. and Ishikawa, S. (2012). Fish Collection Building and Procedures Manual, nglish Edition. Kagoshima: The Kagoshima University Museum.

Nevatte, R. J., Wueringer, B. E., Jacob, D. E., Park, J. M. and Williamson, J. E. (2017). First insights into the function of the sawshark rostrum through examination of rostral tooth microwear. J. Fish Biol. 91, 1582–1602. 10.1111/jfb.13467

Paredes-Aliaga, M. V., Botella, H. and Romero, A. (2024). Testing dental microwear as a proxy for characterising trophic ecology in fossil elasmobranchs (chondrichthyans). Swiss J. Palaeontol. 143, 29. 10.1186/s13358-024-00322-9

Paredes-Aliaga, M. V. and Herráiz, J. L. (2024). Analysing trophic competition in Otodus megalodon and Carcharodon carcharias through 2D-SEM dental microwear. Span. J. Palaeontol. 39, 91–102. 10.7203/sjp.28830

Purnell, M. A., Seehausen, O. and Galis, F. (2012). Quantitative three-dimensional microtextural analyses of tooth wear as a tool for dietary discrimination in fishes. J. R. Soc. Interface 9, 2225–2233. 10.1098/rsif.2012.0140

Purnell, M. A. (1995). Microwear on conodont elements and macrophagy in the first vertebrates. Nature 374, 798–800. 10.1038/374798a0

Purnell, M. A., Bell, M. A., Baines, D. C., Hart, P. J. B. and Travis, M. P. (2007). Correlated evolution and dietary change in fossil stickleback. Science 317, 1887.

Purnell, M. A. and Darras, L. P. G. (2016). 3D tooth microwear texture analysis in fishes as a test of dietary hypotheses of durophagy. Surf. Topogr. 4. 10.1088/2051-672X/4/1/014006

Purnell, M. A., Goodall, R. H., Thomson, S. and Matthews, C. J. (2017). Tooth microwear texture in odontocete whales: variation with tooth characteristics and implications for dietary analysis. Biosurf. Biotribol. 3, 184–195. 10.1016/j.bsbt.2017.11.004

Purnell, M. A., Hart, P. J. B., Baines, D. C. and Bell, M. A. (2006). Quantitative analysis of dental microwear in threespine stickleback: a new approach to analysis of trophic ecology in aquatic vertebrates. J. Anim. Ecol. 75, 967–977. 10.1111/j.1365-2656.2006.01116.x

Purnell, M. A. and Jones, D. (2012). Quantitative analysis of conodont tooth wear and damage as a test of ecological and functional hypotheses. Paleobiology 38, 605–626.

Sallan, L. C. (2014). Major issues in the origins of ray-finned fish (Actinopterygii) biodiversity. Biol. Rev. 89, 950–971. 10.1111/brv.12086

Schmidt, C. W. and Ungar, P. S. (2023). Dental microwear analysis: wear we are going, wear we have been. In A Companion to Biological Anthropology (ed. C. S. Larsen), pp. 572–586. Hoboken, NJ: John Wiley & Sons, Inc.

Standen, E. M., Du, T. Y. and Larsson, H. C. E. (2014). Developmental plasticity and the origin of tetrapods. Nature 513, 54–58. 10.1038/nature13708

Towle, I. and Loch, C. (2021). Tooth chipping prevalence and patterns in extant primates. Am. J. Phys. Anthropol. 175, 292–299. 10.1002/ajpa.24232

Ungar, P. S., Scott, P. S., Scott, J. R. and Teaford, M. (2008). Dental microwear analysis: historical perspectives and new approaches. In Technique and Application in Dental Anthropology (ed. J. D. Irish and G. C. Nelson), pp. 389–425. Cambridge: Cambridge University Press. 10.1017/cbo9780511542442.017

Usami, K. and Kubo, M. O. (2025). In-vitro puncture experiment using alligator teeth tracks the formation of dental microwear and its association with hardness of the diet. Anat. Rec. 10.1002/ar.25659

Vandenplas, S., De Clercq, A. and Huysseune, A. (2014). Tooth replacement without a dental lamina: the search for epithelial stem cells in Polypterus senegalus. J. Exp. Zool. B Mol. Dev. Evol. 322, 281–293. 10.1002/jez.b.22577

Van Valkenburgh, B. (2009). Costs of carnivory: tooth fracture in Pleistocene and Recent carnivorans. Biol. J. Linn. Soc. 96, 68–81. 10.1111/j.1095-8312.2008.01108.x

Van Valkenburgh, B., Peterson, R. O., Smith, D. W., Stahler, D. R. and Vucetich, J. A. (2019). Tooth fracture frequency in gray wolves reflects prey availability. eLife 8, e48628. 10.7554/eLife.48628

Wainwright, P. C., McGee, M. D., Longo, S. J. and Hernandez, L. P. (2015). Origins, innovations, and diversification of suction feeding in vertebrates. Integr. Comp. Biol. 55, 134–145. 10.1093/icb/icv026

Weber, K., Winkler, D. E., Kaiser, T. M., Žigaitė, Ž. and Tütken, T. (2021). Dental microwear texture analysis on extant and extinct sharks: ante-or post-mortem tooth wear? Palaeogeogr. Palaeoclimatol. Palaeoecol. 562, 110147. 10.1016/j.palaeo.2020.110147

Whitlow, Katrina R., Callum F. Ross, Nicholas J. Gidmark, J. D. Laurence-Chasen and Mark W. Westneat. (2022) Suction feeding biomechanics of Polypterus bichir: investigating linkage mechanisms and the contributions of cranial kinesis to oral cavity volume change. Journal of Experimental Biology 225, jeb243283.

Wibisana, J. N., Sallan, R. A., Ota, T., Puchenkov, P., Kubo, T. and Sallan, L. (2025). Modifiable clinical dental impression methods to obtain whole-mouth and detailed dental traits from vertebrates. J. Morphol. 286, e70017. 10.1002/jmor.70017

Winkler, D. E., Iijima, M., Blob, R. W., Kubo, T. and Kubo, M. (2022). Controlled feeding experiments with juvenile alligators reveal microscopic dental wear texture patterns associated with hard-object feeding. Front. Ecol. Evol. 10. 10.3389/FEVO.2022.95772

